# Probing the evolutionary robustness of two repurposed drugs targeting iron uptake in *Pseudomonas aeruginosa*

**DOI:** 10.1101/195974

**Authors:** Chiara Rezzoagli, David Wilson, Michael Weigert, Stefan Wyder, Rolf Kümmerli

## Abstract

**Background and objectives:** Treatments that inhibit the expression or functioning of bacterial virulence factors hold great promise to be both effective and exert weaker selection for resistance than conventional antibiotics. However, the evolutionary robustness argument, based on the idea that anti-virulence treatments disarm rather than kill pathogens, is controversial. Here we probe the evolutionary robustness of two repurposed drugs, gallium and flucytosine, targeting the iron-scavenging pyoverdine of the opportunistic human pathogen *Pseudomonas aeruginosa*.

**Methodology:** We subjected replicated cultures of bacteria to two concentrations of each drug for 20 consecutive days in human serum as an ex-vivo infection model. We screened evolved populations and clones for resistance phenotypes, including the restoration of growth and pyoverdine production, and the evolution of iron uptake by-passing mechanisms. We whole-genome sequenced evolved clones to identify the genetic basis of resistance.

**Results:** We found that mutants resistant against anti-virulence treatments readily arose, but their selective spreading varied between treatments. Flucytosine resistance quickly spread in all populations due to disruptive mutations in *upp*, a gene encoding an enzyme required for flucytosine activation. Conversely, resistance against gallium arose only sporadically, and was based on mutations in transcriptional regulators, upregulating pyocyanin production, a redox-active molecule promoting siderophore-independent iron acquisition. The spread of gallium resistance could be hampered because pyocyanin-mediated iron delivery benefits resistant and susceptible cells alike.

**Conclusions and implications:** Our work highlights that anti-virulence treatments are not evolutionarily robust *per se*. Instead, evolutionary robustness is a relative measure, with specific treatments occupying different positions on a continuous scale.

## Introduction

There is currently much interest in therapeutic approaches that inhibit the expression or functioning of bacterial virulence factors [1–8]. Virulence factors are structures and molecules that allow bacteria to establish and maintain infections [9, 10]. Examples of virulence factors include flagella and pili to adhere to the host tissue, secreted enzymes, tissue-damaging toxins and siderophores to scavenge iron from the host [11]. Approaches that target these traits are called anti-virulence treatments. There is great hope that disarming rather than killing pathogens is an efficient and evolutionarily robust way to manage infections [2, 12–15]. In particular, it is assumed that anti-virulence treatments exert weaker selection for resistance than conventional antibiotics because pathogens are not killed directly. However, empirical evidence for the evolutionary robustness of anti-virulence treatments is controversial with positive and negative reports currently balancing each other out [16–21].

The controversy entails both conceptual and practical aspects. On the conceptual level, some define anti-virulence approaches as treatments that specifically target virulence factors without affecting pathogen growth [2, 22], while others argue that it is unlikely that virulence factors do not affect pathogen fitness, and thus simply use the mechanistic part of the definition [5, 15]. On the practical level, there are debates about what exactly a resistance phenotype is [15], as it could include restoration of virulence factor production, growth (if affected), and/or the activation of a bypassing mechanism, restoring the virulence phenotype [19]. Moreover, there is a shortage of studies examining resistance evolution under realistic conditions in replicated populations, both at the phenotypic and genetic level.

Here, we tackle these issues by examining the mechanistic and evolutionary potential of resistance evolution against two repurposed drugs, gallium and flucytosine, which both target the iron-scavenging pyoverdine of the opportunistic human pathogen *Pseudomonas aeruginosa* [19, 23, 24]. Pyoverdine is an important virulence factor during acute infections [19, 25–31]. It is required to obtain iron from host proteins, such as transferrin and lactoferrin [32]. Given its importance, it has been proposed that drugs interfering with iron uptake could be effective therapeutics to control infections [33]. Gallium and flucytosine both fulfill this role, albeit through different modes of action. Gallium, a repurposed cancer drug, is an iron-mimic and binds irreversibly to secreted pyoverdine, thereby rendering the molecules useless for iron uptake [19, 23, 31]. Flucytosine, a repurposed anti-fungal drug, enters the bacterium, where it is enzymatically activated to a fluorinated ribonucleotide. This active form inhibits, via a yet unknown mechanism, the expression of the *pvdS* iron starvation sigma factor controlling pyoverdine synthesis [24, 34].

In a first set of experiments, we examined whether these two drugs affect the growth of *P. aeruginosa* in human blood serum, a medium that has recently been established as an ex-vivo infection model [35]. We hypothesize that gallium and flucytosine are likely to reduce pathogen fitness as they induce iron starvation [19, 23, 36, 37]. In addition, anti-virulence drugs, like any other drugs, might have deleterious off-target effects affecting growth. Gallium at high dosage, for instance, can penetrate into bacterial cells, where it interferes with redox-active enzymes [38, 39]. Flucytosine, once activated, is known to affect RNA synthesis, which might negatively affect growth [40].

In a second experiment, we examined whether mutants, resistant against these two repurposed drugs, evolve and spread through bacterial populations. To this end, we exposed replicated populations of *P. aeruginosa* to two different concentrations of gallium and flucytosine in human serum. Together with a drug-free control treatment, we let the treated populations evolve for 20 consecutive days in eight-fold replication, by transferring a fraction of the evolving cultures to fresh human serum on a daily basis. Following experimental evolution, we screened evolved populations and clones for possible resistance phenotypes, including the restoration of growth, restoration of virulence factor production, and the evolution of a bypassing mechanism for iron uptake [15, 19]. Finally, we sequenced the whole genome of evolved clones to uncover the genetic basis of potential resistance mechanisms.

Resistance evolution requires two processes: the supply of mutations conferring resistance and appropriate selection regimes favoring the spread of these mutants [41]. With regard to mutation supply, some common resistance mechanisms (e.g. drug degradation, prevention of drug influx and increased drug efflux) are less likely to apply for gallium, which is an ion and acts outside the cell [19]. Therefore, with fewer possible routes to resistance being available, we predict gallium to show higher evolutionarily robustness than flucytosine. However, as for the spread of mutants, both drugs could be evolutionarily robust because they target a secreted virulence factor, which can be shared as a public good between pathogen individuals (iron-loaded pyoverdine can be taken up by all bacteria with a matching receptor; [42, 43]). Consequently, if resistance entails the resumption of virulence factor production then resistant mutants should not spread because they bear the cost of resumed virulence factor production, whilst sharing the benefit with everyone else in the population, including the drug-susceptible individuals [12, 14, 16, 20]. Conversely, if these drugs have deleterious off-target effects, we predict the evolutionary robustness to decline, and accelerated spread of resistance under drug exposure, as for traditional antibiotics.

## Methodology

### Strains and culturing conditions

We used the genetically well-characterized *P. aeruginosa* PAO1 wildtype strain for all experiments. For some assays, we further used a set of knockout mutants in the PAO1 background as control strains (see Supplementary Table S1). Overnight cultures were grown in 8 ml Lysogeny broth (LB) in 50-ml Falcon tubes, incubated at 37°C, 200 rpm for 18 hours.

For all experiments, we washed overnight cultures with 0.8% NaCl solution and adjusted them to OD_600_ = 2.5. Bacteria were further diluted to a final starting of OD_600_ = 2.5 × 10^−3^. All experiments were carried out in human serum, supplemented with HEPES (50 mM) to buffer the medium at physiological pH. Moreover, to impose a standardized iron limitation across experiments, we added the iron chelator human apo-transferrin (100 μg/ml), which is typically present in blood serum at high concentration, and its co-factor NaHCO_3_ (20 mM). We used gallium (GaNO_3_) and flucytosine (5-Fluorocytosine) as anti-bacterials. All chemicals, including human serum, were purchased from Sigma-Aldrich, Switzerland.

### Growth and virulence factor inhibition curves

To assess the extent to which gallium and flucytosine inhibit PAO1 growth and pyoverdine production, we subjected bacterial cultures to a seven-step antibacterial concentration gradient: 0 - 512 µM for GaNO_3_ and 0-140 µg/ml for flucytosine. Overnight cultures of bacteria were grown and diluted as described above and inoculated into 200 μl of human serum on 96-well plates. Plates were incubated at 37 °C in a Tecan Infinite M-200 plate reader (Tecan Group Ltd., Switzerland). We tracked growth by measuring OD at 600 nm and pyoverdine-associated natural fluorescence (excitation: 400 nm, emission: 460 nm) every 15 minutes for 24 hours. Plates were shaken for 15 seconds (3 mm orbital displacement) prior to each reading event.

### Experimental evolution

We exposed wildtype cultures of PAO1 to experimental evolution for 20 days under five different selective regimes in eight-fold replication. The five regimes included a no-drug control, and a low and a high concentration treatment for both drugs (gallium: 50 μM and 280 μM; flucytosine: 10 μg/ml and 140 μg/ml). The antibacterial concentrations were inferred from the dose-response curves (Fig. 1). To initiate experimental evolution, an overnight culture of PAO1 was grown as described above, and individual wells on a 96-well plate were inoculated with 10 μl of culture (diluted to a final density of 10^6^ cells per well) in 190 μl iron-limited human serum. Incubation occurred in the plate reader at 37°C for 23.5 hours, and OD_600_ was measured every 15 minutes prior to a brief shaking event. Subsequently, cultures were diluted in 0.8% NaCl and transferred to a new plate containing fresh media. We adjusted the dilution factor proportional to the overall growth per treatment; no-drug control: 2 × 10^−3^ (day 1-10) and 4*10^−3^ (day 11-20); antibacterial treatments: 10^−3^ (day 1-10) and 2 × 10^−3^ (day 11-20). Following transfers, we added 100 μl of a 50% glycerol-LB solution to cultures for storage at −80°C.

**Figure 1.**
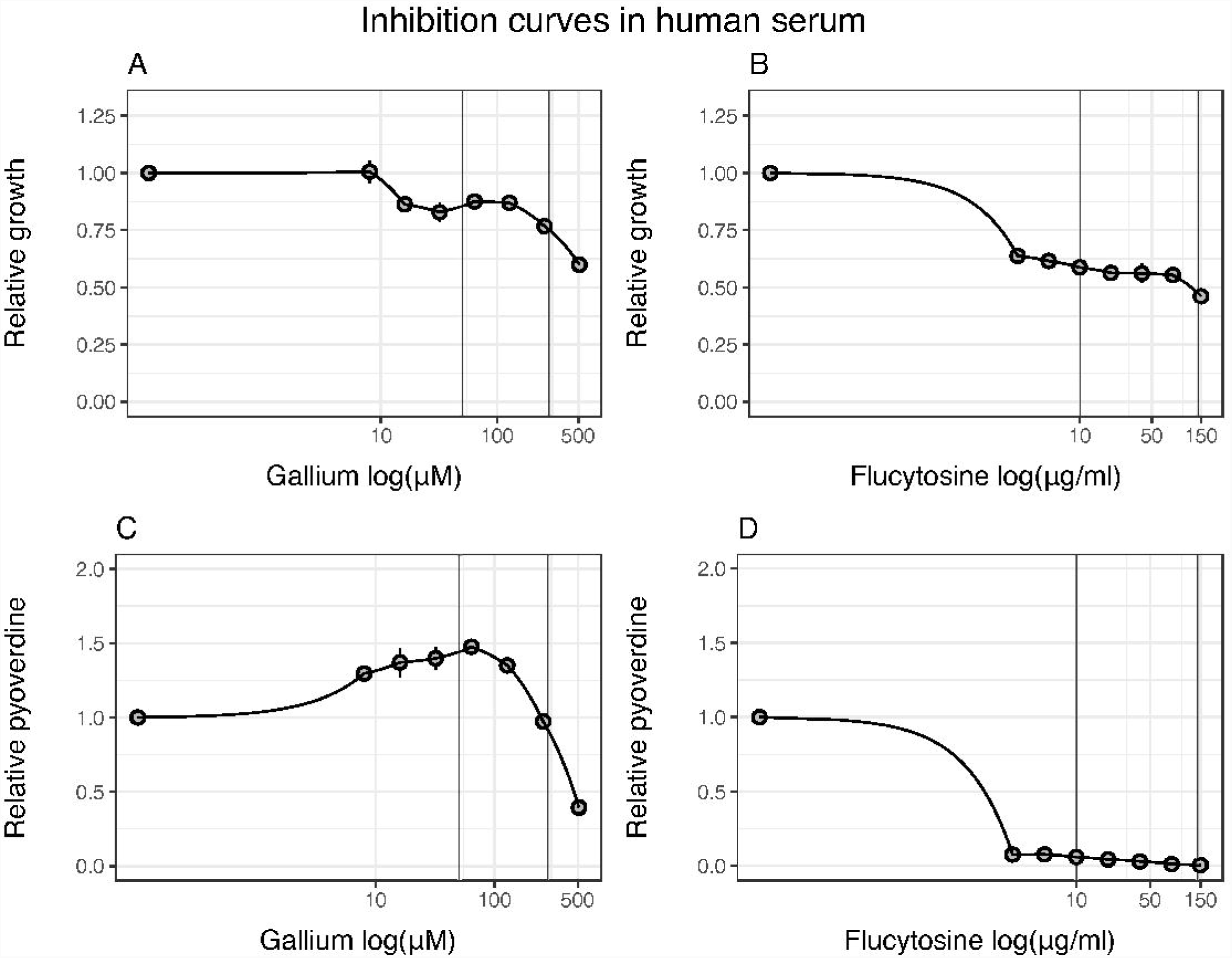
Gallium and flucytosine affect both growth and pyoverdine production of *P. aeruginosa* in human serum (HS). Both anti-virulence drugs reduce growth of bacterial cultures in a dose-dependent manner (A, B), albeit following different patterns: gallium curbs bacterial growth only at relatively high concentrations (A), whereas flucytosine already reduces growth at low concentrations (B). Both drugs further affect pyoverdine production (C, D). When increasing gallium exposure, bacteria first upregulate pyoverdine production at intermediate drug concentrations, but then down-scale investment levels at high drug concentrations (C). In contrast, flucytosine administration leads to an almost complete abolishment of pyoverdine production even at the lowest drug concentration. All data are expressed as average of growth yield, scaled relative to the drug-free treatment. Error bars denote standard errors of the mean across 6 (for flucytosine) and 18 (for gallium) replicates. Dose-response curves were fitted using a spline fit. Vertical lines indicate the drug concentrations used in the experimental evolution.

### Quantification of resistance profiles

To test whether populations evolved under antibacterial exposure restored growth and/or pyoverdine production, we exposed evolved lineages to the drug concentrations they experienced during experimental evolution in 5-fold replication. Following a standard protocol with incubation at 37° C, shaking at 160 rpm, for 24 hours [44], we compared the OD_600_ and pyoverdine-associated fluorescence of evolved lineages relative to the ancestor wildtype grown under drug and no-drug treatment.

To assess potential resistance profiles of individual clones, we streaked out aliquots of evolved lineages onto LB plates. After overnight incubation at 37°C, we randomly picked 200 clones (five colonies per lineage), and assessed their growth and pyoverdine production in 3-fold replication, as described above. Moreover, we performed an in-depth analysis for 20 (four per treatment) randomly picked single clones by quantifying their drug-inhibition curve, following the protocol described above.

To test whether bacteria upregulated alternative iron-acquisition mechanisms, we quantified pyocyanin and protease production of selected clones. For pyocyanin production, overnight bacterial cultures were inoculated into 1 ml of LB (starting OD_600_ = 10^−6^), and incubated at 37°C for 24 hours, shaken at 160 rpm. We measured pyocyanin in the cell-free supernatant through absorbance at 691 nm [19]. For protease production, overnight bacterial cultures were inoculated in human serum (starting OD_600_ = 2.5 × 10^−3^), and incubated at 37°C for 24 hours, shaken at 160 rpm. Subsequently, we centrifuged cultures at 3700 rpm for 15 minutes to obtain protease-containing supernatants. To measure proteolytic activity, we adapted the protocol by [45]: 0.1 ml azocasein solution (30 mg/ml) were mixed with 0.3 ml 50 mM phosphate buffer (pH 7.5), and 0.1 ml culture supernatant. During incubation at 37°C (2 hours), proteases hydrolyze azocasein and release the azo-dye. Proteolytic reaction was stopped by adding 0.5 ml 20% trichloroacetic acid (TCA), samples centrifuged at 12000 rpm (10 min), and proteolytic activity measured through absorbance of the azo-dye at 366 nm.

### Sequencing analysis

We further isolated the genomic DNA of the selected 16 clones evolved under drug regimes and sequenced their genomes. We used the GenElute Bacterial Genomic DNA kit (Sigma Aldrich) for DNA isolation. DNA concentrations were assayed using the Quantifluor dsDNA sample kit (Promega). Samples were sent to the Functional Genomics Center Zurich for library preparation (Nextera XT) and sequencing. Sequencing was performed on the Illumina HiSeq 4000 platform with single-end 125 base pair reads. Adapter sequences were clipped using Trimmomatic v0.33 [46] and reads trimmed using Flexbar v2.5 [47]. We aligned the reads to the PAO1 reference genome using BWA v0.7.12 [48]. We applied GATK v3.5 [49] indel realignment, duplicate removal and HaplotypeCaller SNP/INDEL discovery according to the GATK Best Practices recommendations. This generated a variant call format (VCF) file, from which the following variants were discarded: (i) coverage < 20 reads; (ii) Fisher Strand (FS) score > 30.0, ensuring that there is no strand bias in the data; (iii) QD value < 2.0 (confidence value that there is a true variation at a given site); and (iv) clustered variants (≥ 3 variants in 35nt window) as they likely present sequencing or alignment artifacts. This filtering process yielded a list of potential SNPs and small INDELs, which we annotated using snpEff 4.1g [50] and then screened manually, compared to the sequenced genome of our ancestor wildtype for relevant mutations in gene coding and intergenic regions (Supplementary Table S2).

### Statistical analysis

We used RStudio for statistical analysis (version 0.99.896, with R version 3.3.0). We analyzed growth curves and pyoverdine production profiles using the *grofit* package [51]. We fitted non-parametric model (Splines) curves to estimate growth yield and integral (area under the curve). For all analyses, we scaled growth yield and pyoverdine production relative to the untreated ancestral wildtype. We used general linear mixed effect models to compare whether growth parameters or pyoverdine profiles differ in evolved cultures treated with or without antibacterials. To test for differences between evolved lines and the ancestral wildtype, we used Welch‘s two-sample *t*-test. To compare the dose-response curve of evolved clones, we first fitted spline curves to the inhibition curves, then estimated the integrals of these fits, and compared the scaled fits relative to the ancestor wildtype using ANOVA (Analysis of variance). Protease and pyocyanin production of evolved clones and the ancestor wildtype were corrected for cell number (OD_600_) and analyzed using ANOVA.

## Results

### Gallium and flucytosine curb growth and pyoverdine production in human serum

To confirm that human serum is an iron-limited media, in which pyoverdine is important for growth, we compared the growth of our wildtype strain PAO1 to the pyoverdine-negative mutant PAO1 Δ*pvdD* in either pure human serum or human serum supplemented with transferrin (Supplementary Fig. S1). As expected for iron-limited media, we observed significantly reduced growth of the siderophore-deficient mutants compared to the wildtype (ANOVA: *t*_49_ = −8.13, *p* < 0.0001) under both conditions.

We then subjected PAO1 to a range of drug concentrations in human serum supplemented with transferrin. The resulting dose-response curves revealed that both drugs significantly affected growth and pyoverdine production, albeit following different patterns (Fig. 1). For gallium, growth reduction was moderate at low concentrations, and only became substantial at high concentrations (GaNO_3_ ≥ 256 µM, Fig. 1A). Gallium treatment affected pyoverdine synthesis in a complex way (Fig. 1C), yet consistent with previous findings [19]: at intermediate gallium concentrations, pyoverdine is up-regulated to compensate for the gallium-induced pyoverdine inhibition, and down-regulated at higher concentrations, when pyoverdine-mediated signaling becomes impaired [23]. For flucytosine, already the lowest concentration caused a substantial growth reduction (Fig. 1B) and completely stalled pyoverdine production, with the reduction remaining fairly constant across the concentration gradient (Fig. 1D). We obtained similar response profiles when growing PAO1 in human serum without added transferrin (Supplementary Fig. S2), indicating that transferrin supplementation does not affect the drugs’ mode of actions. For all subsequent experiments, we used human serum with added transferrin to ensure strong iron limitation and to standardize conditions across experiments.

### Do bacteria evolve population-level resistance to antivirulence treatments?

We subjected PAO1 wildtype cultures to experimental evolution both in the absence and presence of gallium and flucytosine (two concentrations each). Eight independent lines per treatment were daily transferred to fresh human serum for a period of 20 days. Subsequently, we assessed whether evolved populations improved growth and/or pyoverdine production levels compared to the treated ancestral wildtype, which could provide first hints of resistance evolution.

For growth (Fig. 2A), we found that evolved lines grew significantly better under drug exposure than the ancestral wildtype (Welch’s t-tests, gallium low (50 µM): *t*_11.9_ = −4.96, *p* = 0.0003; gallium high (280 µM): *t*_13.3_ = −6.48, *p* < 0.0001; flucytosine low (10 µg/ml): *t*_12.2_ = − 5.09, *p* = 0.0002; flucytosine high (140 µg/ml): *t*_7.5_ = −11.79, *p* < 0.0001). Because growth increase could simply reflect adaptation to media components other than drugs, we also analyzed changes in growth performance of the lines evolved without drugs. It turned out that some of the untreated evolved lineages also showed improved growth compared to the ancestral wildtype, but the overall increase across lines was not significant (*t*_9.1_ = −1.61, *p* = 0.1424, Fig. 2A).

**Figure 2.**
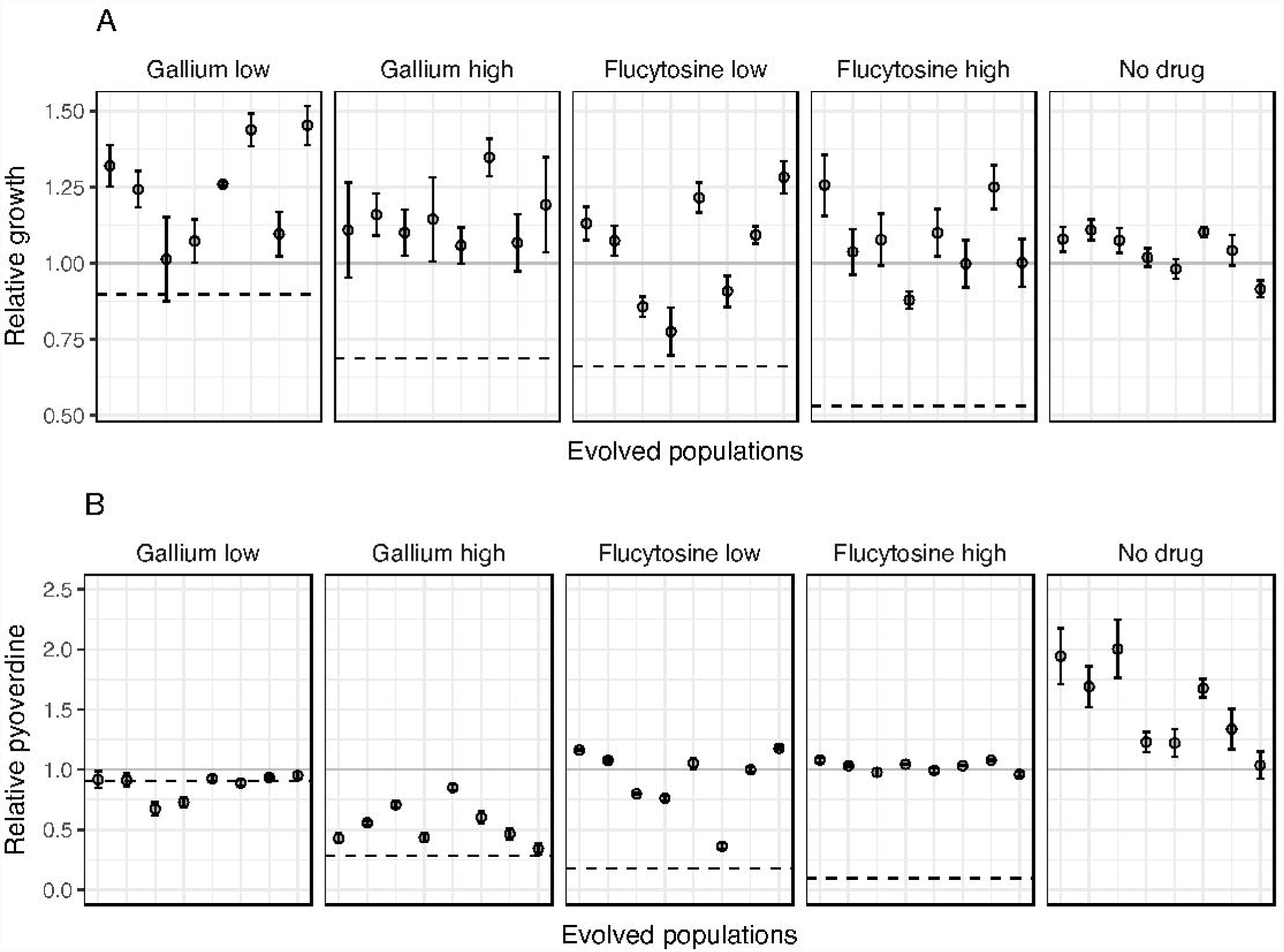
Population level growth and pyoverdine production after evolution in human serum. PAO1 cultures were exposed to either no treatment, low (50 µM) or high (280 µM) gallium, low (10 µg/ml) or high (140 µg/ml) flucytosine concentrations during a 20-day experimental evolution experiment in eight-fold replication. Following evolution, we assessed growth and pyoverdine production of evolved populations (displayed on the x-axis) and compared their performance relative to the untreated (gray solid line, set to 1) and treated ancestral wildtype (black dashed line). (A) Compared to the treated ancestral wildtype, growth of evolved populations significantly increased under all treatment regimes. Growth also increased in some but not all of the non-treatment lines. (B) Pyoverdine production of evolved populations significantly increased relative to the untreated ancestral wildtype under all conditions, also in the no treatment lines. This indicates that increased pyoverdine production might be a general response to growth in human serum, which makes it difficult to disentangle resistance evolution from media adaptation. Error bars show the standard error of the mean across 5 independent replicates.

For pyoverdine production, we observed no significant change for the lines evolved under low gallium concentration (comparison relative to the treated ancestor, Welch’s t-test: *t*_8.8_ = 0.94, *p* = 0.3719) (Fig. 2B). Conversely, lines evolved under the other three drug regimes all showed significantly increased pyoverdine production (Fig. 2B) (gallium high: *t*_13.1_ = −3.69, *p* = 0.0026; flucytosine low: *t*_7.2_ = −7.64, *p* = 0.0001; flucytosine high: *t*_9.6_ = −54.65, *p* < 0.0001). While the increase was moderate for the gallium high treatment, there was full restoration of pyoverdine production in both flucytosine treatments (no significant difference relative to the ancestral untreated wildtype, ANOVA, flucytosine low: *t*_88_ = −1.31, *p* = 0.1944; flucytosine high: *t*_88_ = 0.42, *p* = 0.6766). Although pyoverdine restoration might be taken as evidence for resistance evolution, analysis of the control lines shows that a significant increase in pyoverdine production also occurred in the absence of drugs (Welch’s t-test, *t*_9.1_=−4.03, *p* = 0.0047, Fig. 2B).

### Screening for resistance profiles in evolved single clones

While the population analyses above show that drug resistance and general media adaptation could both contribute to the evolved population growth and pyoverdine phenotypes, we decided to screen individual clones for in-depth analysis. In a first step, we isolated 200 random clones (i.e. 40 per treatment), and individually analyzed their growth and pyoverdine production. These analyses revealed high between-clone variation in growth and pyoverdine production (Supplementary Fig. S3 + S4), suggesting that most evolved populations were heterogeneous, consisting of multiple different genotypes. Interestingly, we observed that 25% of the evolved clones from the no-drug control lines lost the ability to produce pyoverdine (Supplementary Fig. S4). This observation matches the results from previous studies, showing that iron-limitation selects for non-producers that cheat on the pyoverdine produced by others [52, 53]. Increased pyoverdine production at the population level (Fig. 2B) is then typically the result of wildtype cells over-compensating for the presence of non-producers [54, 55]. Conversely, we did not detect non-producers in the four drug treatments, which suggests that selection pressures differ between the non-drug and the drug treatments.

In a second step, we randomly picked 16 single clones (four per drug treatment) and tested whether these evolved clones differ in their drug dose response curve relative to the ancestral wildtype. We observed that three out of eight clones subjected to gallium (Fig. 3A-3D) and all eight clones subjected to flucytosine showed a significantly altered dose response (Fig. 3E-3H). Clones GL_2 and GL_3, evolved under low gallium, showed a significant increase in pyoverdine production under intermediate gallium concentrations (between 8 and 128 μM), which goes along with an improved growth performance for GL_2, but not GL_3. In contrast, clone GH_1, evolved under high gallium concentration, did not show an altered pyoverdine production response, but grew significantly better when exposed to gallium (Fig. 3A-3D). For the eight clones evolved under the flucytosine regime, changes in the dose-response curves were both striking and uniform: growth and pyoverdine production were no longer affected by the drug (Fig. 3E-3H). Since these dose-response curves directly include a control for media adaptation (i.e. the no-drug treatment), our results indicate that all eight clones evolved complete resistance to flucytosine. For gallium, on the other hand, our data suggest that three out of the eight clones exhibited a phenotype that is compatible with at least partial resistance. To check whether these putative resistance profiles are unique to clones evolved under drug treatment, we further assessed the dose-response curves of 4 clones from the no-drug control lines (Supplementary Fig. S5). All these clones responded to both drugs in the same way as the susceptible ancestral wildtype, confirming that adaptation to human serum does not per se result in resistant phenotypes.

**Figure 3.**
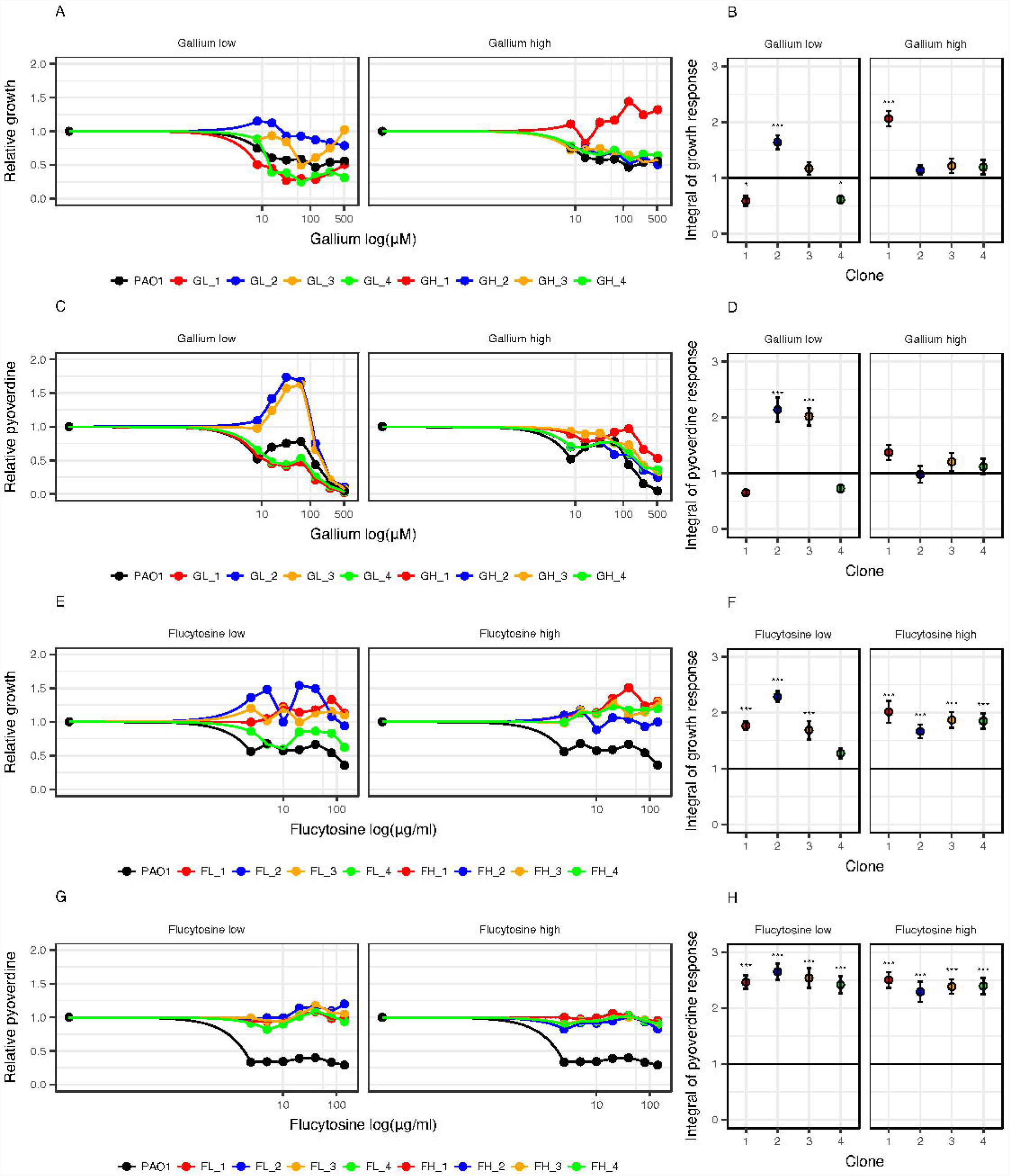
Changes in dose-response curves for evolved single clones indicate resistance evolution. 16 randomly picked clones, four per treatment, were exposed to a range of drug concentrations to test whether their dose-response altered during evolution compared to the ancestral wildtype (black circles and lines). (A and B) Growth dose-response curves under gallium treatment show that two evolved clones (GL2 and GH1) are significantly less inhibited than the ancestral wildtype. (C and D) Pyoverdine dose-response curves under gallium treatment show that two evolved clones (GL2 and GL3) make significantly more pyoverdine than the ancestral wildtype. (E and F) Growth dose-response curves under flucytosine treatment show that all evolved clones grow significantly better than the ancestral wildtype, and are in fact no longer affected by the drug. (G and H) Pyoverdine dose-response curves under flucytosine treatment show that all evolved clones produce significantly more pyoverdine than the ancestral wildtype, and are in fact no longer affected by the drug. Growth and pyoverdine production were measured after 24 h. For each clone, values are scaled relative to its performance in human serum without drugs (absolute values of pyoverdine and growth in the absence of treatment are reported in Supplementary Fig. S6). We used spline functions to fit dose-response curves, and used the integral (area under the curve) to quantify the overall dose response of each clone across the concentration gradient. Error bars denote standard errors of the mean across 6 replicates. Asterisks represent significance levels: * = *p* < 0.05; *** = *p* < 0.0001, based on linear model with *df* = 45.

### Linking phenotypes to genotypes

Our whole-genome sequencing of the 16 focal clones revealed a small number of SNPs and INDELs, which have emerged during experimental evolution (Table 1). All the clones evolved under flucytosine treatment had acquired mutations in the coding sequence of *upp.* There were four different types of mutations, including two different non-synonymous SNPs, a 15-bp deletion and a 1-bp insertion (Supplementary Table S3). The *upp* gene encodes for a uracil phosphoribosyl-transferase, an enzyme required for the intra-cellular activation of flucytosine [56, 57].

**Table 1.**
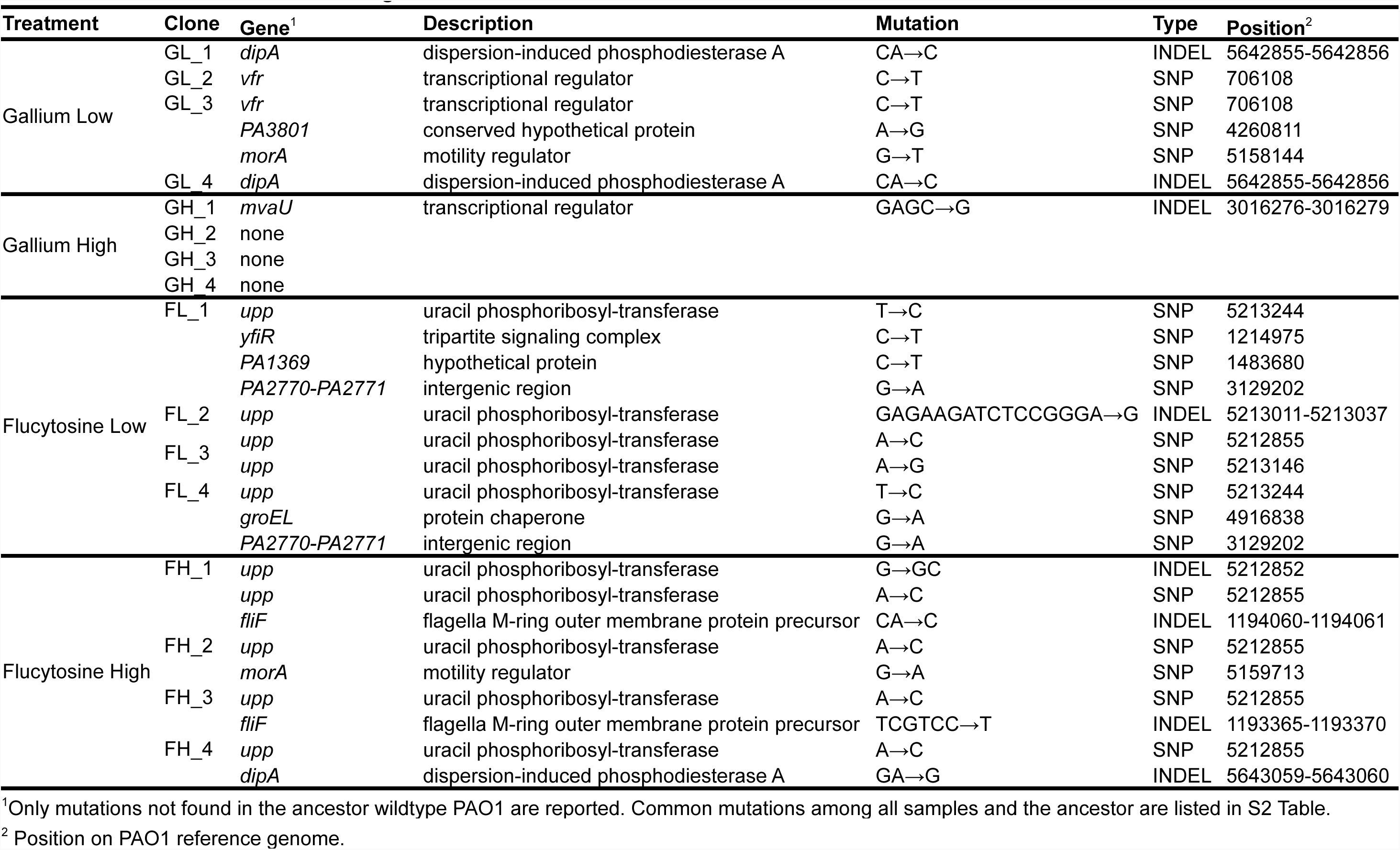
List of mutations in evolved single clones.

For the clones evolved under gallium treatment, the mutational pattern was more heterogeneous (Table 1). No mutations were detected for three clones (GH_2, GH_3, GH_4). In contrast, the three clones with significantly altered dose responses had mutations potentially explaining their phenotypes: clone GH_1 featured a 3-nt deletion in *mvaU*, whereas the clones GL_2 and GL_3 were mutated in *vfr*. Both genes encode transcriptional regulators involved in the regulation of virulence factors, including proteases, pyocyanin and pyoverdine.

In addition, several clones had mutations in *dipA* (dispersion-induced phosphodiesterase A; GL_1, GL_4, FH_4) and *morA* (motility regulator; GL_3, FH_2). The repeated yet unspecific appearance of these mutations could suggest that they represent non-drug-specific adaptations to human serum. Altogether, our sequencing analysis identified three potential targets explaining resistance evolution: the gene *upp* for flucytosine, and the genes encoding the transcriptional regulators *vfr* and *mvaU* for gallium.

### Evolution of bypassing mechanisms for iron acquisition under gallium treatment

It was proposed that bypassing mechanisms, which guarantee iron uptake in a siderophore-independent manner, could confer resistance to gallium [19]. One such by-passing mechanism could involve the up-regulation of pyocyanin, a molecule that can reduce ferric to ferrous iron outside the cell, thereby promoting direct iron uptake [18, 58]. This scenario indeed seems to apply to the three clones mutated in *mvaU* or *vfr*, two regulators that control directly (*mvaU*) or indirectly (*vfr*) the expression of pyocyanin [59, 60]. These clones displayed significantly increased pyocyanin production compared to the ancestral wildtype (Fig. 4A; ANOVA, GH_1: *t*_79_ = 9.64, *p* < 0.0001; GL_2: *t*_99_ = 6.13, *p* < 0.0001; GL_3: *t*_99_ = 14.8, *p* < 0.0001).

**Figure 4.**
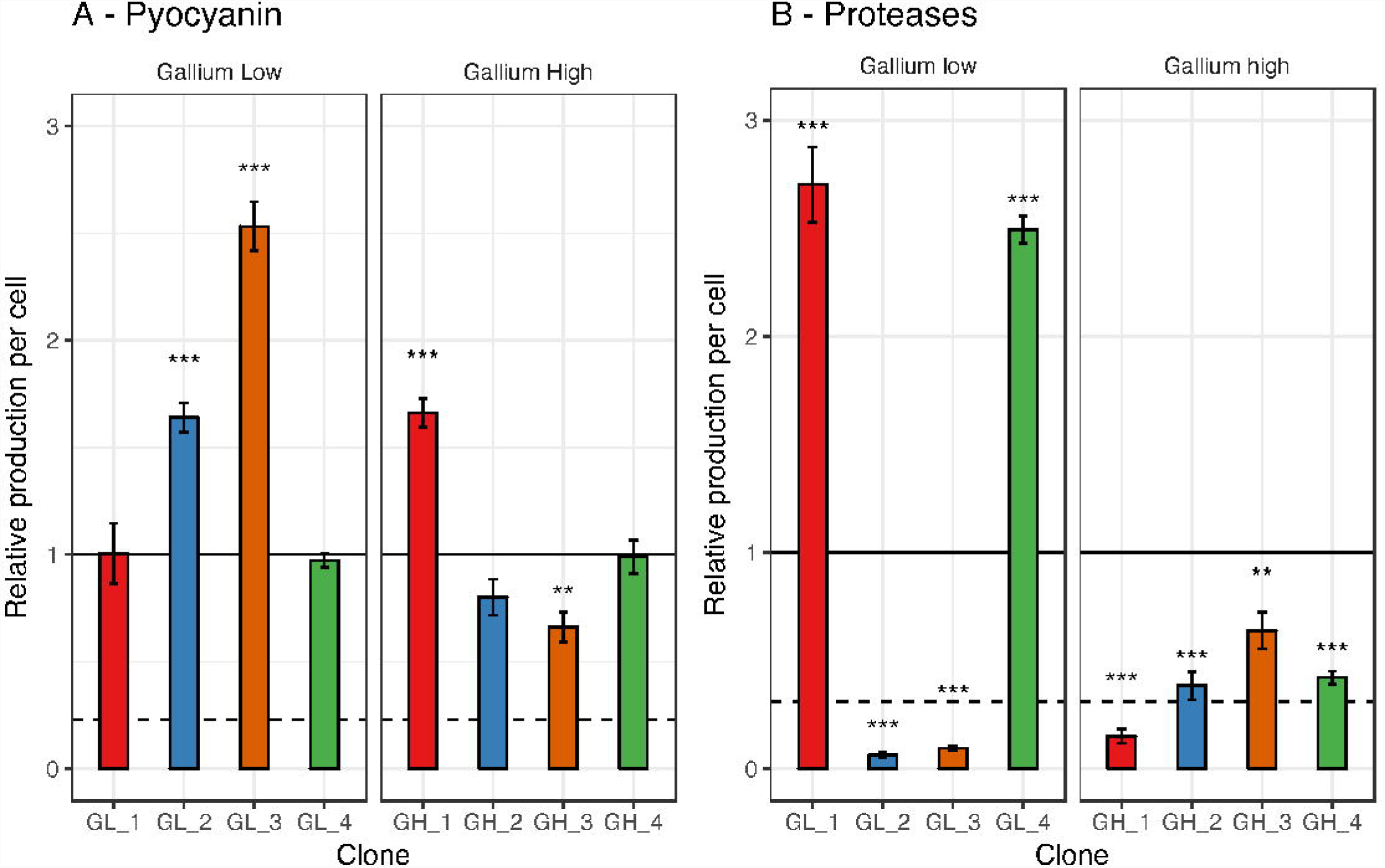
Upregulation of pyocyanin or protease production as potential bypassing mechanisms for iron acquisition under gallium treatment. The eight sequenced clones evolved under gallium treatments (low: 50 µM, high: 280 µM) were screened for their change in the secretion of pyocyanin (A) and proteases (B) relative to the ancestral wildtype. Standard protocols were used for the phenotypic screens in drug free media (see Material and Methods for details). All values are corrected for cell number, and scaled relative to the ancestor wildtype (black line). We included the strains PAO1 Δ*rhlR* (deficient for pyocyanin production) and PAO1 Δ*lasR* (deficient for protease production) as negative controls in the respective assays (dashed lines). Error bars denote standard errors of the mean across 3 (for proteases) and 8 (for pyocyanin) replicates. Asterisks represent significance levels: * = *p* < 0.05; *** = *p* < 0.0001, based on ANOVA.

A second by-passing mechanism could operate via increased protease production, which would allow iron acquisition from transferrin or heme through protease-induced hydrolysis [29, 61]. We found no support for this hypothesis. In fact, six of the evolved clones exhibited reduced and not increased protease activity (Fig. 4B). Moreover, the two clones with significantly increased protease activity (ANOVA, GL_1: *t*_10_ = 13.22, *p* < 0.0001; GL_4: *t*_10_ = 11.60, *p* < 0.0001, Fig. 4B) did not show an altered drug dose-response curve.

### Inactivation of Upp is responsible for resistance to flucytosine

Next, we tested whether the mutations in *upp* are responsible for flucytosine resistance. The enzyme Upp (uracil phosphoribosyl-transferase) is essential for the activation of flucytosine within the cell. The natural function of Upp is to convert uracil to the nucleotide precursor UMP in the salvage pathway of pyrimidine (Fig. 5A). However, *P. aeruginosa* can also produce UMP through the conversion of L-glutamine and L-aspartate [62] (Fig. 5A), suggesting that *upp* is not essential for pyrimidine metabolism. Mutations in this gene could thus prevent flucytosine activation, and confer drug resistance. To test this hypothesis, we compared the flucytosine dose-response curve of the wildtype strain to an isogenic (transposon) mutant (MPAO1Δ*upp*). Consistent with the patterns of the evolved clones (Fig. 3G), we found that MPAO1Δ*upp* was completely insensitive to flucytosine, with neither growth (Fig. 5B) nor pyoverdine production (Fig. 5C) being affected by the drug. These results indicate that *upp* inactivation is a simple and efficient mechanism to become flucytosine resistant.

**Figure 5.**
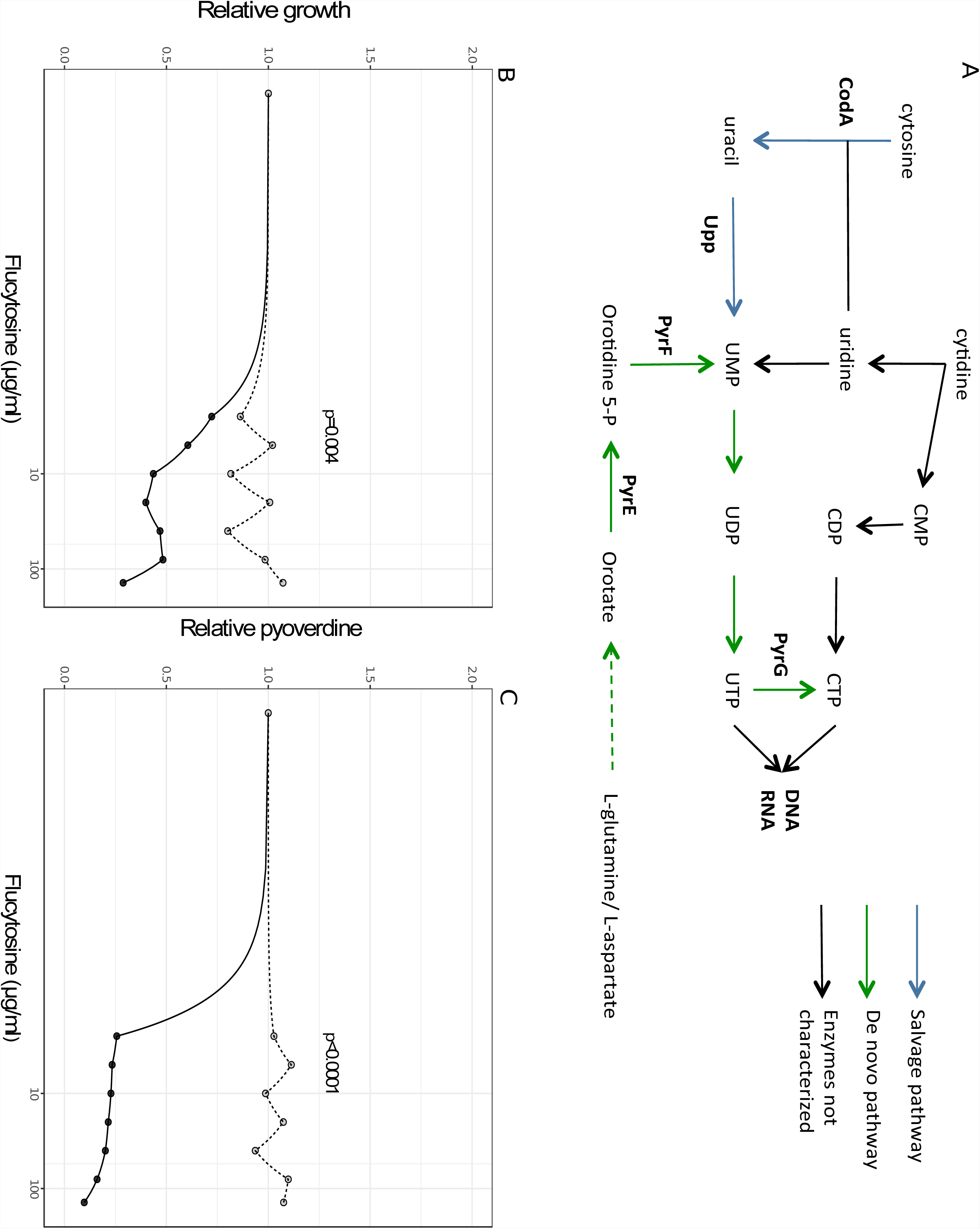
Upp is a non-essential enzyme, and mutations in its gene result in flucytosine resistance. (A) Flucytosine interferes with the pyrimidine metabolism in *P. aeruginosa.* The drug enters the cell through the transporter CodB (not shown), where it is first converted to fluorouracil by the cytosine deaminase CodA, and then to fluoro-UMP by the uracil phosphoribosyl-transferase Upp. Fluoro-UMP is a modified nucleotide precursor, the action of which results in RNA molecules with compromised functionality. Through an as yet unknown mechanism, fluoro-UMP also arrests pyoverdine synthesis in *P. aeruginosa*. Importantly, the nucleotide-precursor UMP can also be produced through an alternative *de-novo* pathway from the amino acids L-glutamine and L-aspartate, making Upp a non-essential enzyme in this bacterium. Experiments with the transposon mutant MPAO1 Δ*upp* (deficient for Upp production) indeed demonstrate that the lack of Upp no longer affects strain growth (B) and pyoverdine production (C). This demonstrates that the inactivation of *upp* is a simple and efficient way to evolve resistance to flucytosine. Experiments were carried out in human serum across a range of flucytosine concentrations. Growth and pyoverdine production of MPAO1 Δ*upp* (gray-dashed lines) and its corresponding wildtype MPAO1 (black-solid lines) were measured after 24 hours for each treatment separately in 6-fold replication. All values are scaled relative to the drug-free treatment. For statistical analysis, we compared the integrals of the dose-response curves between the mutant and the wildtype strain (Welch’s *t*-test, growth: *t*_4.3_ = −5.56, *p* = 0.0041; pyoverdine production: *t*_5.0_ = −48.80, *p* < 0.0001).

## Discussion

New treatment approaches against the multi-drug resistant ESKAPE pathogens, to which *P. aeruginosa* belongs, are desperately needed [8, 63, 64]. In this context, treatments that disarm rather than kill bacteria have attracted particular interest, because such approaches have been proposed to be both effective in managing infections and sustainable in the sense that resistance should not easily evolve [2, 5, 12–15]. Promising approaches include the quenching of toxins [65, 66], siderophores required for iron-scavenging [19, 23, 24, 37, 67], and quorum sensing molecules regulating virulence factor production [3]. In our study, we probed the evolutionary robustness argument by focusing on two repurposed drugs (gallium and flucytosine) targeting siderophore production of *P. aeruginosa*. Using a combination of replicated experimental evolution and phenotypic and genotypic analysis, we show that the often-recited argument of anti-virulence drugs being evolutionarily robust is not supported. Instead, we provide a nuanced view on the molecular mechanisms and selective forces that can lead to resistance. For flucytosine, for instance, we found repeated resistance evolution based on a mechanism that prevents drug activation inside the cell, which mitigates possible pleiotropic and deleterious effects caused by this drug. For gallium, meanwhile, two types of partially resistant mutants, based on siderophore bypassing mechanisms, arose. However, these mutants only sporadically emerged, indicating that their potential to selective spread in populations is compromised. Our work highlights that evolutionary robustness is a relative measure with specific treatments lying on different positions on a continuum. Thus, our task is not to argue about whether anti-virulence drugs are evolutionarily robust or not, but to assess the relative position of each novel treatment on this continuum.

Our findings indicate that it is difficult to define anti-virulence treatments based on fitness effects [2, 6, 8, 22]. This is because fitness effects might vary in response to the ecological context of the media or the infection. For instance, prior work [24] showed that flucytosine does not affect bacterial growth in trypticase soy broth dialysate (TSBD), whereas we found significant fitness effects in human serum. Endorsing the fitness-based definition would mean that flucytosine could only be considered as an anti-virulence drug in very specific cases, i.e. in one media but not in another. This might cause confusion when we aim to bring these new treatment approaches to the clinic. While we agree that it would be ideal to find compounds that only curb virulence but not fitness, it seems that such cases are rare and context-dependent [5, 15]. For all those reasons, we support the more general definition of antivirulence treatments as advocated in previous reviews [5, 15]: drugs intended to target bacterial virulence factors.

Important to note is that even when we use the more general definition of anti-virulence the chances are good that many of the new treatment approaches are evolutionary more robust than classical antibiotics. This is nicely illustrated in the case of gallium, where we found that partially resistant mutants only sporadically occurred. Given the mutation rate in *P. aeruginosa* and the number of generations that occurred in our experiment, the frequency of such mutants should be much higher if they had experienced a clear selective advantage (Table 2). Our data thus highlight that it is important to distinguish between the appearance of resistant mutants and their evolutionary potential to spread through populations. At the mechanistic level, we isolated mutants with increased pyocyanin production, a potential mechanism to by-pass gallium-mediated pyoverdine quenching. Pyocyanin is a redox active molecule that can extracellularly reduce ferric to ferrous iron [18, 58]. The upregulation of pyocyanin was associated with mutations in *mvaU*, encoding a positive regulator of pyocyanin production, and *vfr*, encoding a global virulence factor regulator [10]. Mutations in Vfr can activate PQS (Pseudomonas Quinolon Signal) synthesis, which is known to promote pyocyanin and pyoverdine synthesis [60, 68]. At the evolutionary level, however, the selective advantage of these mutations seemed to be compromised because they occurred only in some of the sequenced clones (Table 1). One plausible explanation for their sporadic appearance is that pyocyanin could serve as a public good, reducing iron outside the cell, thereby generating benefits for other individuals in the vicinity, including the drug-susceptible wildtype cells. This scenario would support the argument that anti-virulence strategies should target collective traits, because this would prevent resistant mutants to fix in populations [12, 14, 16, 19, 20]. The relative success of these mutants is then determined by the viscosity of the environment, determining the shareability of secreted compounds [69], and the potential for negative-frequency dependent selection, where strain frequency settles at an intermediate ratio [70–72].

**Table 2.**
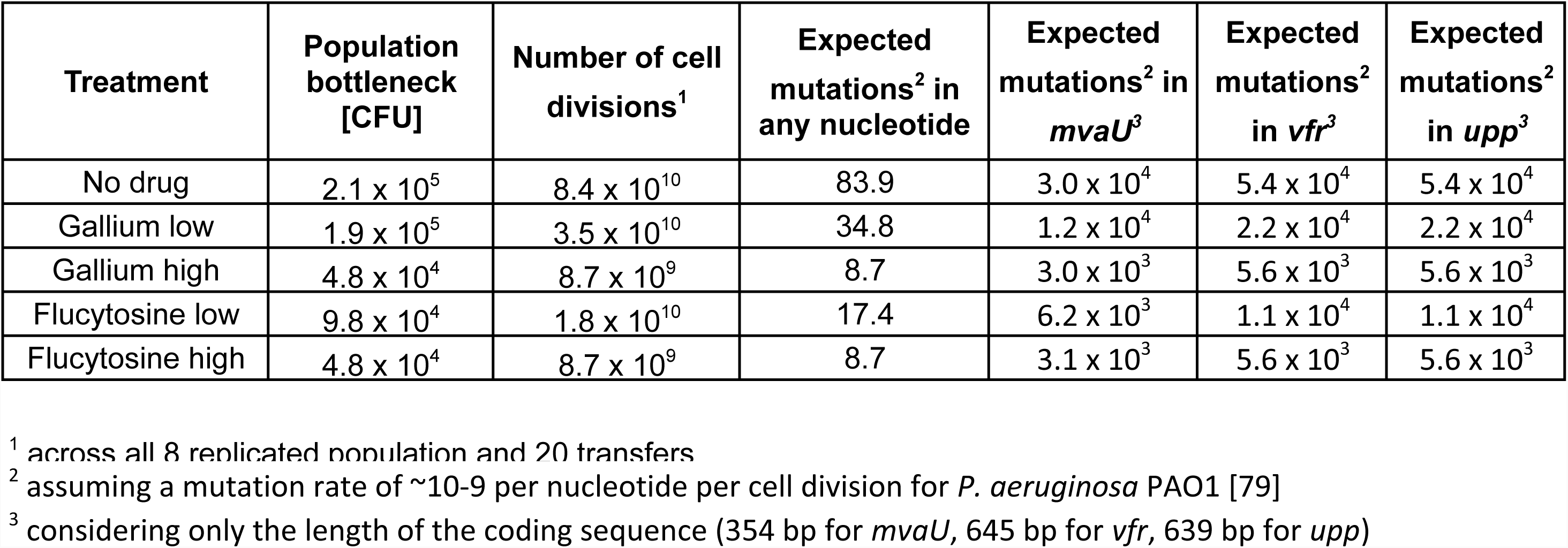
Estimation of mutation supply during experimental evolution.

The pattern clearly differed for flucytosine, where we found pervasive resistance evolution. Although it is not exactly known how flucytosine inhibits pyoverdine synthesis, we argue that resistance evolution could mainly be caused by negative effects on other traits than pyoverdine synthesis. Flucytosine undergoes several enzymatic modifications within the cell, finally resulting in fluorinated ribonucleotides. While flucytosine was shown to inhibit pyoverdine synthesis [24], it likely also interferes with nucleotide synthesis, which might compromise RNA functionality more generally [73]. This sets the stage for selection to favor mutants with alleviated fitness costs under drug exposure. Our results suggest that cells achieved this through mutations in *upp*. The scheme depicted in Fig. 5A shows that the essential pyrimidine nucleotide precursor UMP can be synthesized either through the salvage pathways reutilizing exogenous free bases and nucleosides, or via a *de novo* biosynthesis pathway using L-glutamine or L-aspartate. While the salvage pathway is typically preferred because it requires less energy, it generates the harmful fluoro-UMP under flucytosine treatment. Thus, the abolishment of the salvage pathway through mutations in *upp* and the switching to the *de novo* biosynthesis pathway provides a selective advantage under flucytosine exposure. The notion that off-target effects might compromise the evolutionary robustness of anti-virulence drugs, is also supported by the work of Maeda *et al.* [17]. They showed that resistance to the quorum-quenching compound C-30 (brominated furanone) evolves repeatedly via upregulation of a drug efflux pump. The spread of these mutants in their experiment can be explained by the fact that quorum quenching did not only inhibit virulence factor production, but also compromised the ability of cells to grow in adenosine medium, which requires a functional quorum sensing system [74].

## Conclusions and Implications

Our work advances research on anti-virulence drugs on multiple fronts. First, it shows that resistant phenotypes are difficult to define, as they can involve the restoration of growth, the resumption of virulence factor production, and/or the activation of a bypassing mechanism. Detailed phenotypic and genotypic analyses, as those proposed in our study, are required to disentangle background adaptation from resistance evolution. Second, we show that anti-virulence approaches are neither completely evolution-proof nor does the notion “all roads lead to resistance” apply [75]. A detailed evolutionary analysis for each individual drug is required to assess its position on the continuum between the two extremes. Third, we advocate the application of more rigorous evolutionary approaches to quantify resistance evolution. While there are rigorous standards to describe the precise molecular mode of action of a novel antibacterial [21, 76], there is much room for improvement for standards regarding the quantification and characterization of resistance evolution [77, 78].

## Acknowledgments

We thank Adin Ross-Gillespie for advice, the Functional Genomics Center Zurich for technical support with the strain sequencing.

## Supplementary data

Supplementary data are available at EMPH online. Whole-genome sequencing data have been deposited in the ArrayExpress database at EMBL-EBI (www.ebi.ac.uk/arrayexpress) under accession number E-MTAB-6110.

## Supporting information captions

**Supplementary Figure S1. Human Serum is an iron limited media.** To show that pyoverdine production is beneficial in human serum, we compared growth of *P. aeruginosa* PAO1 wildtype and the siderophore-deficient mutant PAO1 Δ*pvdD*, in pure human serum or human serum supplemented with the strong iron chelator human apo-transferrin (100 μg/ml) and its co-factor NaHCO_3_ (20 mM). When adding only 50 mM HEPES to buffer the medium at physiological pH (first panel) we observed that growth of the siderophore mutant was significantly reduced compared to the wildtype (Welch’s t-test, *t*_9_= 7.31, *p* < 0.0001), confirming that human serum is an iron-limited media. When we increased iron limitation (second panel), we found that overall growth decreased compared to pure human serum (ANOVA, growth in human serum + transferrin: *t*_21_= −22.35, *p* < 0.0001), but that the wildtype PAO1 still grew significantly better compared to the siderophore mutant. (Welch’s t-test, *t*_10_= 3.30, *p* < 0.0086), confirming that the siderophore pyoverdine is important for iron scavenging in human serum. All growth data are scaled relative to the wildtype growth in human serum supplemented with HEPES 50 mM. Error bars denote standard errors of the mean across 6 replicates.

**Supplementary Figure S2. Gallium and flucytosine affect growth and pyoverdine production of *P. aeruginosa* in plain human serum, not supplemented with additional transferrin.** To test the possibility that the two drugs affect growth and pyoverdine differently in human serum with and without additional transferrin, we performed dose-response curves in plain human serum, without adding the iron-chelator. We found that both anti-virulence drugs reduce growth of bacterial cultures (A, B) and pyoverdine production (C, D) in a dose-dependent manner, similarly to the reduction observed in human serum with transferrin (Fig. 1). As PAO1 has higher growth potential in plain serum (Supplementary Fig. S1A), the inhibitory effect of the two drugs at high concentrations was more pronounced than in serum with transferrin. Error bars denote standard errors of the mean across 6 replicates. Dose-response curves were fitted using a spline fit.

**Supplementary Figure S3. Growth and pyoverdine production profiles of evolved single clones under treatment regimes.** We were interested in examining the variation in growth and pyoverdine production profiles among evolved clones. For that purpose, we streaked out all evolved cultures on LB agar, and picked 5 random clones per evolved population. Overall, we isolated 160 clones, 40 clones per treatment. Single clones were tested under the treatment regime experienced during experimental evolution, and growth and pyoverdine production were measured after 24 hours. Panels show growth and pyoverdine production of evolved clones relative to the untreated wildtype (grey line) under gallium (A, C) and flucytosine treatment (B, D). We found heterogeneous growth and pyoverdine patterns under all treatment regimes, suggesting diversification in evolving populations. Data points show means across three independent replicates. Dashed lines depict the mean growth or the mean pyoverdine production of the ancestral strain under treatment. Labeled single clones marked in red refer to the clones used for in-depth analysis and sequencing.

**Supplementary Figure S4. Pyoverdine non-producer strains evolved in human serum with transferrin in the no-drug control treatment.** The strong iron limitation in human serum with transferrin could alone exert selective pressure during the experimental evolution, independently from the drug. This could affect the ability of the evolved clones to produce siderophore in response to the iron-limitation. To investigate these effects and control for general adaptation to low iron conditions, we isolated 40 clones from each treatment (gallium low, gallium high, flucytosine low; flucytosine high; no-drug control) and screened their ability to produce pyoverdine in human serum with transferrin. We observed that in the populations evolved without drug, pyoverdine-negative clones evolved at a frequency of approx. 25%. On the contrary, all clones from the drug treatments still produced pyoverdine. Each data point represents a single measurement per evolved clone. The black line denotes the average wildtype production level in the same assay and the grey area refers to the wildtype mean ± standard error of 20 replicates.

**Supplementary Figure S5: Evolved clones from the no-drug control treatment remain sensitive to both gallium and flucytosine.** We were interested in determining whether media adaptation could per se lead to reduced susceptibility to gallium and flucytosine. To test this, we picked 4 random clones from the no-drug evolved population and subjected them to a range of drug concentration, for both gallium and flucytosine, to measure if they still respond to the drug. Growth dose-response curves under treatment showed that all evolved clones were equally or more sensitive to both gallium (A-B) and flucytosine (E-F), compared to the ancestor PAO1. Similarly, pyoverdine dose-response curves showed that all clones were still affected by gallium (C-D). Under flucytosine, three clones, were slightly less sensitive to flucytosine compared to the ancestor (G-H), although they can still considered sensitive to the antimicrobial if compared to the resistant clones isolated from the flucytosine high or low treatment (Figure 3 G-H). Growth and pyoverdine production were measured after 24 h. For each clone, values are scaled relative to its performance in human serum without drugs. We used spline functions to fit dose-response curves, and used the integral (area under the curve) to quantify the overall dose response of each clone across the concentration gradient. Error bars denote standard errors of the mean across 6 replicates. Asterisks represent significance levels: * = p < 0.05; *** = p < 0.0001, based on linear model with df = 30.

**Supplementary Figure S6. Growth and pyoverdine production of evolved single clones in human serum with no treatment.** We quantified the growth (A) and pyoverdine production (B) of the 16 evolved single clones we used for in-depth analysis and whole-genome sequencing. Specifically, we grew the clones for 24 hours in human serum and compared their performance to the ancestor wildtype in absence of the treatments. This allowed us to test for media adaptation. (A) We observed that some of the clones from both treatments showed significantly improved growth compared to the ancestral wildtype. This was the case for clones GL1 (*t*_43_ = 2.04, *p* = 0.0475), GL_4 (*t*_43_ = 2.04, *p* = 0.0475), GH_1 (*t*_43_ = 3.13, *p* = 0.0125), FL_1 (*t*_43_ = 5.88, *p* < 0.0001) and all clones evolved under high flucytosine (FH_1: *t*_43_ = 3.09, *p* = 0.0054; FH_2:, *t*_43_ = 3.03, *p* = 0.0054; FH_3: *t*_43_ = 4.38, *p* = 0.0002; FH_4, *t*_43_ = 2.82, *p* = 0.0071). Interestingly, two of the clones evolved under gallium low treatment showed reduced growth compared to PAO1 (GL_2: *t*_43_ = −3.35, *p* = 0.0033; GL_3: *t*_43_ = −3.36, *p* = 0.0033). (B) Regarding per capita pyoverdine production (pyoverdine fluorescence divided by growth), we found that two clones showed increased levels of pyoverdine production under gallium treatment (GL_1: *t*_43_ = 2.79, *p* = 0.0200; GL_4: *t*_43_ = 2.69, *p* = 0.0200), and three clones of the flucytosine high treatment did so too (FH_1: t_43_ = 2.43, p = 0.0356; FH_2: *t*_43_ = 2.29, *p* = 0.0356; for FH_4: *t*_43_ = 2.35, *p* = 0.0356). In contrast, there were also a number clones with significantly reduced pyoverdine production compared to PAO1 (GH_1: *t*_43_= −3.14, *p* = 0.0045; GH_3: *t*_43_= −3.10, *p* = 0.0045; GH_4: *t*_43_= −3.46, *p* = 0.0045; FL_4: *t*_43_= −2.02, *p*=0.0488). All data are scaled relative to the ancestor wildtype. Error bars denote standard error of the mean across 6 replicates. Asterisks represent significance codes (* = p<0.05; ** = p<0.001, *** = p<0.0001 based on ANOVA, corrected for multiple pairwise comparison with the false discovery rate method).

**Supplementary Figure S7. Alteration of the fluorescent properties of pyoverdine when bound to gallium.** The fluorescent signal of pyoverdine becomes inflated when gallium binds to it [19]. To take this signal bias into account, we quantified the bias in fluorescence signal as a function of gallium concentration. Specifically, we supplemented iron-limited human serum with 200 μM of purified pyoverdine across a range of gallium concentrations (8 to 512 μM, as used for the main experiments). We found that the signal bias can be explained by a 3 parameters-logistic function. We used this function to correct for signal bias in all our analyses.

**Supplementary Table S1. List of control strains used in this study.**

**Supplementary Table S2. Differences to the PAO1 reference genome shared among all sequenced evolved clones and the ancestor wildtype.**

**Supplementary Table S3. Effects of mutation on the Upp protein sequence of evolved single clones**

## References

1. Escaich S. Antivirulence as a new antibacterial approach for chemotherapy. Curr Opin Chem Biol 2008;12:400–8.

2. Rasko DA, Sperandio V. Anti-virulence strategies to combat bacteria-mediated disease. Nat Rev Drug Discov 2010;9:117–28.

3. LaSarre B, Federle MJ. Exploiting Quorum Sensing To Confuse Bacterial Pathogens. Microbiol Mol Biol Rev 2013;77:73–111.

4. Maura D, Ballok AE, Rahme LG. Considerations and caveats in anti-virulence drug development. Curr Opin Microbiol 2016;33:41–6.

5. Vale PF, McNally L, Doeschl-Wilson A et al. Beyond killing. Evol Med Public Heal 2016;2016:148–57.

6. Johnson BK, Abramovitch RB. Small Molecules That Sabotage Bacterial Virulence. Trends Pharmacol Sci 2017;38:339–62.

7. Rampioni G, Visca P, Leoni L et al. Drug repurposing for antivirulence therapy against opportunistic bacterial pathogens. Emerg Top Life Sci 2017:ETLS20160018.

8. Dickey SW, Cheung GYC, Otto M. Different drugs for bad bugs: antivirulence strategies in the age of antibiotic resistance. Nat Rev Drug Discov 2017;16:1–15.

9. Rahme LG, Stevens EJ, Wolfort SF et al. Common virulence factors for bacterial pathogenicity in plants and animals. Science (80-) 1995;268:1899–902.

10. Balasubramanian D, Schneper L, Kumari H et al. A dynamic and intricate regulatory network determines Pseudomonas aeruginosa virulence. Nucleic Acids Res 2013;41:1–20.

11. Wu HJ, Wang AHJ, Jennings MP. Discovery of virulence factors of pathogenic bacteria. Curr Opin Chem Biol 2008;12:93–101.

12. André JB, Godelle B. Multicellular organization in bacteria as a target for drug therapy. Ecol Lett 2005;8:800–10.

13. Baron C. Antivirulence drugs to target bacterial secretion systems. Curr Opin Microbiol 2010;13:100–5.

14. Pepper JW. Drugs that target pathogen public goods are robust against evolved drug resistance. Evol Appl 2012;5:757–61.

15. Allen RC, Popat R, Diggle SP et al. Targeting virulence: can we make evolution-proof drugs? Nat Rev Microbiol 2014;12:300–8.

16. Mellbye B, Schuster M. The sociomicrobiology of antivirulence drug resistance: a proof of concept. MBio 2011;2:e00131–11.

17. Maeda T, García-Contreras R, Pu M et al. Quorum quenching quandary: resistance to antivirulence compounds. ISME J 2012;6:493–501.

18. García-Contreras R, Lira-Silva E, Jasso-Chávez R et al. Isolation and characterization of gallium resistant Pseudomonas aeruginosa mutants. Int J Med Microbiol 2013;303:574–82.

19. Ross-Gillespie A, Weigert M, Brown SP et al. Gallium-mediated siderophore quenching as an evolutionarily robust antibacterial treatment. Evol Med Public Heal 2014;2014:18–29.

20. Gerdt JP, Blackwell HE. Competition Studies Confirm Two Major Barriers That Can Preclude the Spread of Resistance to Quorum-Sensing Inhibitors in Bacteria. ACS Chem Biol 2014;9:2291–9.

21. Sully EK, Malachowa N, Elmore BO et al. Selective Chemical Inhibition of agr Quorum Sensing in Staphylococcus aureus Promotes Host Defense with Minimal Impact on Resistance. PLoS Pathog 2014;10:e1004174.

22. Clatworthy AE, Pierson E, Hung DT. Targeting virulence: a new paradigm for antimicrobial therapy. Nat Chem Biol 2007;3:541–8.

23. Kaneko Y, Thoendel M, Olakanmi O et al. The transition metal gallium disrupts Pseudomonas aeruginosa iron metabolism and has antimicrobial and antibiofilm activity. J Clin Invest 2007;117:877–88.

24. Imperi F, Massai F, Facchini M et al. Repurposing the antimycotic drug flucytosine for suppression of Pseudomonas aeruginosa pathogenicity. Proc Natl Acad Sci U S A 2013;110:7458–63.

25. Meyer JM, Neely A, Stintzi A et al. Pyoverdin is essential for virulence of Pseudomonas aeruginosa. Infect Immun 1996;64:518–23.

26. Takase H, Nitanai H, Hoshino K et al. Impact of siderophore production on Pseudomonas aeruginosa infections in immunosuppressed mice. Infect Immun 2000;68:1834–9.

27. Harrison F, Browning LE, Vos M et al. Cooperation and virulence in acute Pseudomonas aeruginosa infections. BMC Biol 2006;4:21.

28. Cornelis P, Dingemans J. Pseudomonas aeruginosa adapts its iron uptake strategies in function of the type of infections. Front Cell Infect Microbiol 2013;3:1–7.

29. Bonchi C, Imperi F, Minandri F et al. Repurposing of gallium-based drugs for antibacterial therapy. BioFactors 2014;40:303–12.

30. Granato ET, Harrison F, Kümmerli R et al. Do Bacterial “Virulence Factors” Always Increase Virulence? A Meta-Analysis of Pyoverdine Production in Pseudomonas aeruginosa As a Test Case. Front Microbiol 2016;7:1–13.

31. Weigert M, Ross-Gillespie A, Leinweber A et al. Manipulating virulence factor availability can have complex consequences for infections. Evol Appl 2017;10:91–101.

32. Valenti P, Berlutti F, Conte MP et al. Lactoferrin Functions: Current Status and Perspectives. J Clin Gastroenterol 2004;38:S127–9.

33. Smith DJ, Lamont IL, Anderson GJ et al. Targeting iron uptake to control Pseudomonas aeruginosa infections in cystic fibrosis. Eur Respir J 2013;42:1723–36.

34. Visca P, Imperi F, Lamont IL. Pyoverdine siderophores: from biogenesis to biosignificance. Trends Microbiol 2007;15:22–30.

35. Bonchi C, Frangipani E, Imperi F et al. Pyoverdine and proteases affect the response of Pseudomonas aeruginosa to gallium in human serum. Antimicrob Agents Chemother 2015;59:5641–6.

36. Banin E, Lozinski A, Brady KM et al. The potential of desferrioxamine-gallium as an antiPseudomonas therapeutic agent. Proc Natl Acad Sci U S A 2008;105:16761–6.

37. DeLeon K, Balldin F, Watters C et al. Gallium Maltolate Treatment Eradicates Pseudomonas aeruginosa Infection in Thermally Injured Mice. Antimicrob Agents Chemother 2009;53:1331–7.

38. Chitambar CR. Gallium-containing anticancer compounds. Future Med Chem 2012;4:1257–72.

39. Hijazi S, Visca P, Frangipani E. Gallium-Protoporphyrin IX Inhibits Pseudomonas aeruginosa Growth by Targeting Cytochromes. Front Cell Infect Microbiol 2017;7:1–15.

40. Waldorf AR, Polak A. Mechanisms of action of 5-fluorocytosine. Antimicrob Agents Chemother 1983;23:79–85.

41. Hughes D, Andersson DI. Evolutionary Trajectories to Antibiotic Resistance. Annu Rev Microbiol 2017;71:579–96.

42. Griffin AS, West SA, Buckling A. Cooperation and competition in pathogenic bacteria. Nature 2004;430:1024–7.

43. Inglis RF, Biernaskie JM, Gardner A et al. Presence of a loner strain maintains cooperation and diversity in well-mixed bacterial communities. Proc R Soc B Biol Sci 2016;283, doi:10.1098/rspb.2015.2682.

44. Kümmerli R, Jiricny N, Clarke LS et al. Phenotypic plasticity of a cooperative behaviour in bacteria. J Evol Biol 2009;22:589–98.

45. Chessa JP, Petrescu I, Bentahir M et al. Purification, physico-chemical characterization and sequence of a heat labile alkaline metalloprotease isolated from a psychrophilic Pseudomonas species. Biochim Biophys Acta - Protein Struct Mol Enzymol 2000;1479:265–74.

46. Bolger AM, Lohse M, Usadel B. Trimmomatic: a flexible trimmer for Illumina sequence data. Bioinformatics 2014;30:2114–20.

47. Dodt M, Roehr J, Ahmed R et al. FLEXBAR—Flexible Barcode and Adapter Processing for Next-Generation Sequencing Platforms. Biology (Basel) 2012;1:895–905.

48. Li H, Durbin R. Fast and accurate short read alignment with Burrows–Wheeler transform. Bioinformatics 2009;25:1754–60.

49. McKenna A, Hanna M, Banks E et al. The Genome Analysis Toolkit: a MapReduce framework for analyzing next-generation DNA sequencing data. Genome Res 2010;20:1297–303.

50. Cingolani P, Platts A, Wang LL et al. A program for annotating and predicting the effects of single nucleotide polymorphisms, SnpEff. Fly (Austin) 2012;6:80–92.

51. Kahm M, Hasenbrink G, Ludwig J. grofit: Fitting Biological Growth Curves with R. J Stat Softw 2010;33:1–21.

52. Harrison F, Paul J, Massey RC et al. Interspecific competition and siderophore-mediated cooperation in Pseudomonas aeruginosa. ISME J 2008;2:49–55.

53. Dumas Z, Kümmerli R. Cost of cooperation rules selection for cheats in bacterial metapopulations. J Evol Biol 2012;25:473–84.

54. Harrison F. Dynamic social behaviour in a bacterium: Pseudomonas aeruginosa partially compensates for siderophore loss to cheats. J Evol Biol 2013;26:1370–8.

55. Ross-Gillespie A, Dumas Z, Kümmerli R. Evolutionary dynamics of interlinked public goods traits: An experimental study of siderophore production in Pseudomonas aeruginosa. J Evol Biol 2015;28:29–39.

56. Beck DA, O’Donovan GA. Pathways of pyrimidine salvage in Pseudomonas and former Pseudomonas: Detection of recycling enzymes using high-performance liquid chromatography. Curr Microbiol 2008;56:162–7.

57. Edlind TD, Katiyar SK. Mutational Analysis of Flucytosine Resistance in Candida glabrata. Antimicrob Agents Chemother 2010;54:4733–8.

58. Cox CD. Role of pyocyanin in the acquisition of iron from transferrin. Infect Immun 1986;52:263–70.

59. Li C, Wally H, Miller SJ et al. The multifaceted proteins MvaT and MvaU, members of the H-NS family, control arginine metabolism, pyocyanin synthesis, and prophage activation in Pseudomonas aeruginosa PAO1. J Bacteriol 2009;191:6211–8.

60. Diggle SP, Matthijs S, Wright VJ et al. The Pseudomonas aeruginosa 4-Quinolone Signal Molecules HHQ and PQS Play Multifunctional Roles in Quorum Sensing and Iron Entrapment. Chem Biol 2007;14:87–96.

61. Doring G, Pfestorf M, Botzenhart K et al. Impact of Proteases on Iron Uptake of Pseudomonas aeruginosa Pyoverdin from Transferrin and Lactoferrin. Infect Immun 1988;56:291–3.

62. Isaac JH, Holloway BW. Control of pyrimidine biosynthesis in Pseudomonas aeruginosa. J Bacteriol 1968;96:1732–41.

63. Pendleton JN, Gorman SP, Gilmore BF. Clinical relevance of the ESKAPE pathogens. Expert Rev Anti Infect Ther 2013;11:297–308.

64. Brown D. Antibiotic resistance breakers: can repurposed drugs fill the antibiotic discovery void? Nat Rev Drug Discov 2015;14:821–32.

65. Lu C, Maurer CK, Kirsch B et al. Overcoming the Unexpected Functional Inversion of a PqsR Antagonist in Pseudomonas aeruginosaLJ: An In Vivo Potent Antivirulence Agent Targeting pqs Quorum Sensing. Angew Chemie Int Ed 2014;53:1109–12.

66. Henry BD, Neill DR, Becker KA et al. Engineered liposomes sequester bacterial exotoxins and protect from severe invasive infections in mice. Nat Biotechnol 2015;33:81–8.

67. Kelson AB, Carnevali M, Truong-Le V. Gallium-based anti-infectives: targeting microbial iron-uptake mechanisms. Curr Opin Pharmacol 2013;13:707–16.

68. Lin J, Zhang W, Cheng J et al. A Pseudomonas T6SS effector recruits PQS-containing outer membrane vesicles for iron acquisition. Nat Commun 2017;8:14888.

69. Weigert M, Kümmerli R. The physical boundaries of public goods cooperation between surface-attached bacterial cells. Proc R Soc B Biol Sci 2017;284.

70. Ross-Gillespie A, Gardner A, West SA et al. Frequency dependence and cooperation: theory and a test with bacteria. Am Nat 2007;170:331–42.

71. Raymond B, West SA, Griffin AS et al. The dynamics of cooperative bacterial virulence in the field. Science (80-) 2012;337:85–8.

72. Yurtsev EA, Chao HX, Datta MS et al. Bacterial cheating drives the population dynamics of cooperative antibiotic resistance plasmids. Mol Syst Biol 2013;9:683.

73. Harbers E, Chaudhuri NK, Heidelberger C. Studies on fluorinated pyrimidines. VIII. Further biochemical and metabolic investigations. J Biol Chem 1959;234:1255–62.

74. Dandekar AA, Chugani S, Greenberg EP. Bacterial Quorum Sensing and Metabolic Incentives to Cooperate. Science 2012;338:264–6.

75. Breidenstein EBM, de la Fuente-Núñez C, Hancock REW. Pseudomonas aeruginosa: all roads lead to resistance. Trends Microbiol 2011;19:419–26.

76. Ling LL, Schneider T, Peoples AJ et al. A new antibiotic kills pathogens without detectable resistance. Nature 2015;517:455–9.

77. Perron GG, Zasloff M, Bell G. Experimental evolution of resistance to an antimicrobial peptide. Proc R Soc B Biol Sci 2006;273:251 LP–256.

78. Hochberg ME, Jansen G. Bacteria: Assessing resistance to new antibiotics. Nature 2015;519:158.

79. McElroy KE, Hui JGK, Woo JKK, et al. Strain-specific parallel evolution drives short-term diversification during Pseudomonas aeruginosa biofilm formation. Proc Natl Acad Sci 2014;111: E1419–E1427.

